# RNA-dependent structures of the RNA-binding loop in the flavivirus NS3 helicase

**DOI:** 10.1101/2020.01.15.907725

**Authors:** Russell B. Davidson, Josie Hendrix, Brian J. Geiss, Martin McCullagh

## Abstract

The flavivirus NS3 protein is a helicase that has pivotal functions during the viral genome replication process, where it unwinds double-stranded RNA and translocates along the nucleic acid polymer in a nucleoside triphosphate hydrolysis-dependent mechanism. An increased interest in this enzyme as a potential target for development of antiviral therapeutics was sparked by the 2015 Zika virus epidemic in the Americas. Crystallographic and computational studies of the flavivirus NS3 helicase have identified the RNA-binding loop as an interesting structural element, which may function as an origin for the RNA-enhanced NTPase activity observed for this family of helicases. Microsecond-long unbiased molecular dynamics as well as extensive replica exchange umbrella sampling simulations of the Zika NS3 helicase have been performed to investigate the RNA-dependence of this loop’s structural conformations. Specifically, the effect of the bound single-stranded RNA (ssRNA) oligomer on the putative “open” and “closed” conformations of this loop are studied. In the Apo substrate state, the two structures are nearly isoergonic (Δ*G*_*O*→*C*_ = −0.22 kcal mol^−1^), explaining the structural ambiguity observed in Apo NS3h crystal structures. The bound ssRNA is seen to stabilize the “open” conformation (Δ*G*_*O*→*C*_ = 1.97 kcal mol^−1^) through direct protein-RNA interactions at the top of the loop. Interestingly, a small ssRNA oligomer bound over 13 Å away from the loop is seen to affect the free energy surface to favor the “open” structure while minimizing barriers between the two states. The mechanism of the transition between “open” and “closed” states is characterized as are residues of importance for the RNA-binding loop structures. From these results, the loop is hypothesized to be a viable region in the protein for targeted small-molecule inhibition and mutagenesis studies, where stabilization of the “closed” RNA-binding loop will negatively impact RNA-binding and the RNA-enhanced NTPase activity.

## Introduction

The flaviviruses (family *Flaviviridae*) present major health threats to the tropical and subtropical regions of the world. Over half of the world’s population lives in areas that are susceptible to infections from this viral family, which includes dengue, Zika, and West Nile viruses.^1,2^ Outbreaks of Zika virus (ZIKV) in 2013 and 2014, in French Polynesia, as well as 2015, in Brazil, have been correlated with severe congenital malformations (microcephaly) as well as neurological complications such as Guillain-Barré syndrome.^3^ The most recent ZIKV epidemic in the Americas sparked major research endeavors into identification of novel antiviral drugs against Zika and other flaviviruses.^3–7^ Towards this goal, fundamental research on the viral enzymes’ structures and functions will aid the identification of antiviral targets.

The nonstructural protein 3 (NS3) of flaviviruses has been identified as one such target due to its pivotal role during the viral replication cycle. ^7–18^ As one of eight nonstructural proteins encoded by flaviviruses, NS3 is a multifunctional enzyme that possess an N-terminal serine protease domain and a C-terminal helicase/nucleoside triphosphatase (NTPase)/RNA triphosphatase domain.^19–24^ The N-terminal protease is responsible for cleaving the viral polyprotein during translation. The C-terminal helicase domain (NS3h) unwinds the double-stranded RNA replication intermediate during the viral genome replication process. ^25^ In order to do so, NS3h binds and translocates along a single stranded RNA (ssRNA) polymer in a nucleoside triphosphate (NTP) hydrolysis-dependent mechanism. Therefore, the NS3h enzyme represents a complex molecular machine that has pivotal functions in the replication of the viral genome.

NS3h is categorized as a superfamily 2 (SF2) helicase (NS3/NPH-II subfamily) that hydrolyzes NTP to unwind double stranded RNA and translocate along the nucleic acid oligomer in a 3′ to 5′ direction.^26^ A structural representation of ZIKV NS3h is shown in Fig 1A. The enzyme has three subdomains; subdomains 1 and 2 are RecA-like structures with large *β*-sheets forming the core of the subdomain while subdomain 3 is unique to the NS3 subfamily and less conserved structurally across the *Flaviviridae* viruses. The RNA-binding cleft, highlighted in orange, is a large channel that separates subdomain 3 from subdomains 1 and 2, with the 5′ terminus of the ssRNA oligomer positioned at the top of the protein in Fig 1A. The NTP hydrolysis active site, highlighted in purple, is positioned between the beta sheets of subdomains 1 and 2.

**Fig 1.**
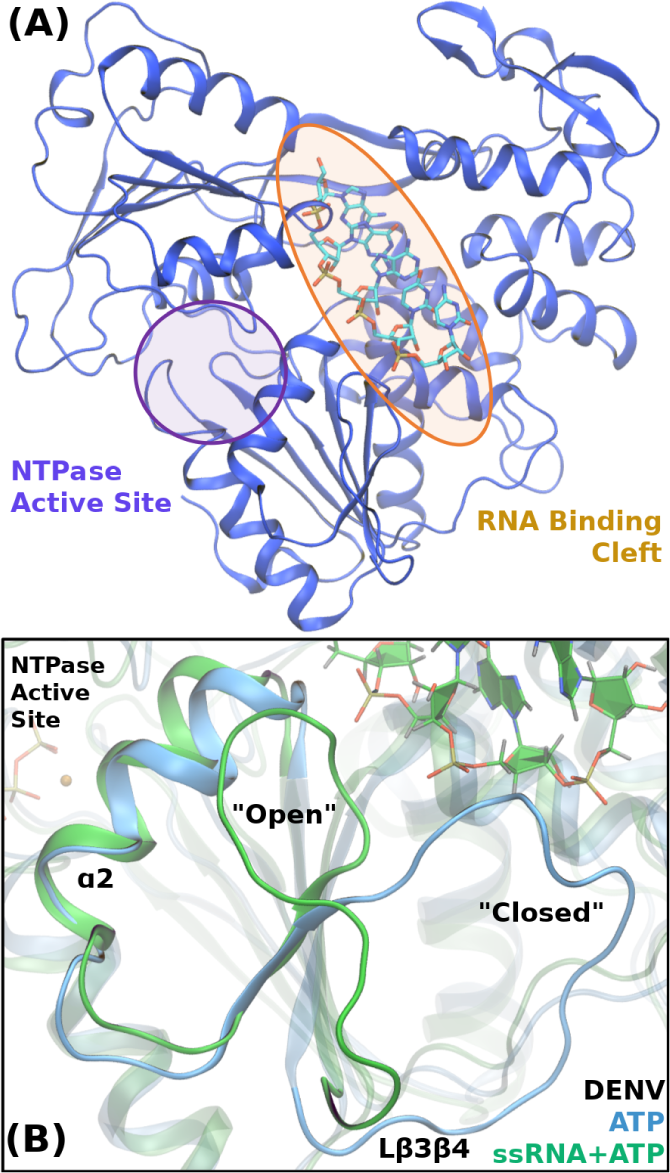
The ssRNA substrate state of the Zika NS3h (PDB 5GJB). The NTPase and RNA-binding clefts are highlighted in purple and orange, respectively. The 3′ terminus of the single-stranded RNA interacts with subdomain 1 and predominantly the RNA-binding loop. This loop region, also named L*β*3*β*4, is depicted in panel B, which has been adapted from Davidson *et al.*^27^ *RNA-induced “open” and “closed” conformations of Lβ*3*β*4 are depicted using the dengue NS3h ATP (blue) and ssRNA+ATP (green) crystal structures due to lack of the well-resolved “closed” conformations in Zika crystal structures.

Both active sites and respective functions are strongly coupled; NS3h is known to exhibit RNA-stimulated NTPase activity and NTPase-dependent helicase activity.^19–24,28^ Coupling between these two active sites has been the subject of numerous recent studies.^27–32^ The RNA-binding loop, shown in Fig 1B, is highlighted in this body of research as a site of allosteric structural change induced by the RNA oligomer, which we hypothesize is one of the origins of the RNA-enhanced NTPase activity.^27^ This loop structure represents a large region of the RNA-binding cleft where the 3′ terminus of the bound RNA interacts with subdomain 1. Positioned between *β*3 and *β*4 of the subdomain 1 *β*-sheet, the RNA-binding loop (also termed L*β*3*β*4) is seen to have two major conformational states in flavivirus NS3h crystal structures. The dengue NS3h crystal structures presented in Fig 1B depict these two states: the “open” conformation has the loop closely interacting with *α*2 (termed the spring helix by Gu and Rice^33^) while, in the “closed” conformation, the loop is positioned in close proximity to *α*3 and is blocking the lower portion of the RNA-binding cleft.

During development of a dengue virus vaccine (DENVax), mutation of a NS3h residue in the RNA-binding loop (E250A) has been identified as one of three mutations producing a significantly attenuated phenotype.^17,18^ It is unclear from these vaccine development studies why such a mutation results in the observed phenotype, specifically an increased temperature sensitivity and decreased viral replication. Yet, these results suggest that slight modification to the RNA-binding loop negatively affects the viral replication process.

The body of crystal structures available on the Protein Data Bank (PDB) of ZIKV and other flavivirus NS3h has left ambiguity as to the functional importance of the conformational states of the RNA-binding loop. Recently, Jain *et al.* highlighted the L*β*3*β*4 structure in their Apo ZIKV NS3h crystal structure (PDB ID: 5JRZ) relative to structures seen in other flavivirus NS3h.^29^ In this crystal structure, the RNA-binding loop was resolved in the “open” conformation, which was insightful because the “open” conformation had previously been associated with RNA-bound structures.^34,35^ In non-RNA-bound conformations, L*β*3*β*4 had always been resolved in the “closed” conformation or poorly resolved due to high flexibility of the loop region.

Additionally, a pair of molecular dynamics studies have recently provided insight into the L*β*3*β*4 structural states. Mottin *et al.* reported multiple 100 ns simulations of the ZIKV Apo and ssRNA-bound structures.^31^ Large structural fluctuations of L*β*3*β*4 were observed in the Apo simulations with dampened conformational fluctuations seen in the ssRNA simulations. Minimal quantification of the observed L*β*3*β*4 structural states were performed for these simulations. In Davidson *et al.*, we reported micro-second long simulations of dengue NS3h structures where the “open” conformation maintained direct interactions with residues in *α*2, which is seen to push the helix in towards the NTPase active site and, thereby, affect the behaviors of active site water molecules.^27^ These results led us to hypothesize that the RNA-binding loop is an allosteric site where RNA affects the NTPase activity through the loop-*α*2 interactions observed in the “open” conformation.

We report here a directed study of the L*β*3*β*4 conformational states in an effort to understand the effect RNA has on the loop. Micro-second long MD simulations of the ZIKV Apo, ssRNA, and an artificially-altered ssRNA:NS3h systems were used to sample the loop conformations in different RNA-bound substrate states. In the simulation of the Apo system, a transition from the “open” to “closed” conformation is observed and quantified using numerous methods. Finally, replica exchange umbrella sampling (REUS) simulations were used to enhance the sampling of the RNA-binding loop’s conformations in the three substrate states, allowing us to quantify the RNA-induced effect on the loop’s structural free energy landscape. This body of simulations demonstrates that an RNA oligomer perturbs the L*β*3*β*4 free energy surface to favor the “open” conformation over the “closed” conformation, even when the RNA is far removed from the loop.

## Methods

### Starting Structures and System Preparation

Initial structures for the reported all-atom, explicit solvent MD simulations originated from the ZIKV Apo NS3h (PDB: 5JRZ) and binary NS3h:ssRNA (PDB: 5GJB) crystal structures.^29,34^ Additionally, a ssRNA-bound system was artificially created from the 5GJB conformation, where three of the five nucleotides were removed from the 3′ end of the RNA oligomer. The remaining RNA in this structure is positioned at the top of the RNA-binding cleft, ∼ 13.5 Å away from the closest L*β*3*β*4 residue. This structure and respective simulations will be referred to as ssRNA_1−2_.

### Molecular Dynamics Simulations

All-atom, explicit solvent MD simulations were performed for the three conformations discussed above (denoted Apo, ssRNA, and ssRNA_1−2_). The simulations were performed using the GPU-enabled AMBER16 and AMBER18 software, ^36,37^ ff14SB^38^ parameters for proteins, and ff99bsc0_*χ* OL3_ ^39,40^ parameters for RNA. For each system, the starting structures were solvated in TIP3P water boxes with at least a 12 Å buffer between the protein and periodic images. Crystallographic waters were maintained. Sodium and chloride ions were added to neutralize charge and maintain a 0.10 M ionic concentration. The Langevin dynamics thermostat and Monte Carlo barostat were used to maintain the systems at 300 K and 1 bar. Direct nonbonding interactions were calculated up to a 12 Å distance cutoff. The SHAKE algorithm was used to constrain covalent bonds that include hydrogen.^41^ The particle-mesh Ewald method was used to account for long-ranged electrostatic interactions. ^42^ A 2 fs integration time step was used with energies and positions written every 5 ps. The minimum amount of simulation performed for each system was one trajectory of 1.0 *µ*s, with the first 200 ns of simulation sacrificed to equilibration of the starting structures. Simulation of the Apo system was performed to 1.3 *µ*s in order to thoroughly sample the “closed” L*β*3*β*4 conformation.

### Adaptive Sampling of L*β*3*β*4 Transition

Neither the ssRNA nor ssRNA_1−2_ systems sampled the L*β*3*β*4 conformation transition during the respective microsecond MD simulations. To obtain structures of the “closed” conformation, three independent, 100 ns steered molecular dynamics (SMD) simulations were used to slowly pull L*β*3*β*4 into poorly sampled structural space.^43^ For these SMD trajectories, the pulling collective variable was the distance between the Val250 (L*β*3*β*4) and Arg269 (*α*3) C*α* atoms. A pulling force constant of 20 kcal mol^−1^ Å^−2^ was used; all other simulation parameters were maintained as above. The rate of pulling was approximately 0.5 Å ns^−1^.

Frames from this set of SMD trajectories were subsequently used to initiate 40 independent, unbiased trajectories, each 50 ns long. Selection of starting frames for these trajectories was performed using a two dimensional projection of the unbiased and SMD trajectories onto essential dynamics (discussed in the Supplementary information, SI) eigenvectors. Regions of poor sampling in this projection space were identified and frames associated with these regions were used in these independent, unbiased trajectories. This body of simulation represents an extra 2 *µ*s of simulation for both the ssRNA and ssRNA_1−2_ systems.

### Replica Exchange Umbrella Sampling (REUS) Simulations

The REUS method was used to enhance the sampling of the L*β*3*β*4 conformational space for the Apo, ssRNA, and ssRNA_1−2_.^44^ The distance between Ala230 (*α*2) and Ala247 (L*β*3*β*4) C*α* atoms was used as the biased collective variable. This single atomic displacement distance was chosen because it has the largest free energy barrier from the Apo unbiased simulation’s sampling of the “open” to “closed” conformational change. Additionally, the Ala230 C*α* atom has small positional variance due to being in *α*2; the large changes in this distance collective variable occurring during the transition are strongly correlated with the L*β*3*β*4 transition.

For each of the three systems described above, 40 windows were used in the REUS simulations. Equilibrium wells of these windows range from 3.625 Å to 18.250 Å with these equilibrium wells separated by 0.375 Å. Harmonic, biasing force constants were 20 kcal mol^−1^ Å^−2^. Window exchanges were attempted every 25 ps, with an average accepted rate of 0.37 attempts. The biased CV values are written every 200 fs, while frames and energies are written every 2 ps. For each window, a total of 46 ns of REUS simulations was performed, amounting to a total of 1.84 *µ*s of enhanced sampling of the L*β*3*β*4 structures for each of the systems.

Due to the protocol used to obtain structures of the “closed” conformation in the ssRNA and ssRNA_1−2_ systems, the first 10 ns of the REUS simulations were not included in free energy analyses. We use the eigenvector method for umbrella sampling (EMUS) analysis package to calculate the stitched free energy surfaces for these REUS simulations.^45^

### Data Analysis

Analyses of MD trajectories were performed using Python 3.7.4 and the MDAnalysis module (version 0.19.2).^46^ Matplotlib was used for plotting data.^47^ VMD was used for visualization of trajectories and production of structural figures.^48–50^ All analysis scripts are available on Github (https://github.com/mccullaghlab/ZIKV-Lb3b4). Additional analysis details are provided in the SI document.

## Results and Discussion

The structure-function relationship of the RNA-binding loop is poorly understood, especially when considering the RNA-dependence of the loop’s structure. 56 monomeric flavivirus NS3h structures have been reported on the Protein Data Bank (PDB), of which a large proportion have poorly resolved electron densities for L*β*3*β*4 atoms. Additionally, a multiple sequence alignment of the flavivirus NS3h sequences indicates that the loop is one of the least conserved regions of the protein, which generally suggests low functional importance of sequence positions. See the SI document for further discussion of the structural and bioinformatic analyses of flavivirus NS3h. Yet, as presented in Fig 1B, large structural changes are seen in the loop when comparing RNA-bound and Apo structures, a common trend across all flaviviruses. This ambiguity around the RNA-binding loop’s functional importance has motivated our molecular dynamics study of the loop and, more specifically, its RNA-dependent conformational states.

Results of this study are presented in two sections. The first section highlights the L*β*3*β*4 structural states observed during the unbiased, microsecond-long MD simulations of Apo, ssRNA, and ssRNA_1−2_ systems. Focus is given to the thorough quantification of a transition between the “open” and “closed” loop conformational states, observed during the Apo NS3h simulation, as well as specific residues hypothesized to be functionally important during this transition. The second section reports free energy surfaces of the structural transition, as obtained from REUS enhanced sampling simulations, to highlight the effect that the bound RNA oligomer has on L*β*3*β*4’s structure.

### L*β*3*β*4 conformations in MD simulations

Microsecond-long simulations of Zika NS3h Apo (5JRZ), ssRNA (5GJB), and an artificially-altered ssRNA system (ssRNA_1−2_) have been performed to sample the RNA-binding loop’s conformations in the presence or absence of an RNA oligomer. Starting structures for all three systems have L*β*3*β*4 in the “open” conformation, directly interacting with residues of *α*2. Shown in Fig 2A, the root mean square deviation (RMSD) of the loop backbone atoms for these simulations, referenced against the 5GJB structure, indicate that large structural deviations occur during the Apo simulation while RNA-bound systems maintain the “open” loop conformation. The small structural deviations seen during the ssRNA simulation are expected due to strong interactions between the 3′ RNA nucleotides and protein residues at the top of the loop. Surprisingly, artificial removal of three 3′ nucleotides has a limited effect on the L*β*3*β*4 conformational sampling, as seen in Fig 2. This suggests that an RNA-binding loop conformational change is a rare event that may be affected via indirect RNA-loop interactions.

**Fig 2.**
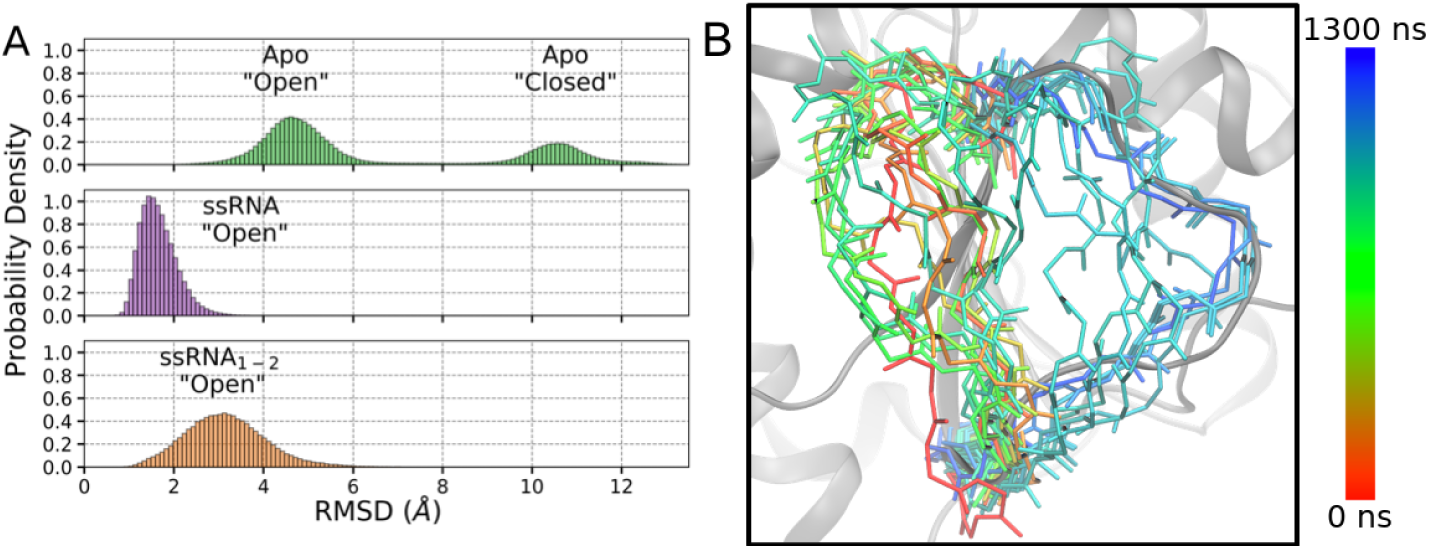
Structural fluctuations of L*β*3*β*4 during the MD simulations. (A) Root mean square deviation (RMSD) analysis of residues 246 to 254, relative to the 5GJB crystal structure. Large RMSD values, seen in the Apo results, indicate a large structural shift away from the “open” loop conformation occurring during that simulation. (B) Structural representation of L*β*3*β*4 backbone atoms over many timesteps of the Apo simulation. At ∼ 900 ns (light green to turquoise), the loop structure begins to transition from the “open” to “closed” conformation.

Large structural deviations of the loop have been observed by Mottin *et al.* in 100 ns MD simulations of the 5JRZ and 5JMT crystal structures.^31^ A large transition between the “open” and “closed” loop conformations is observed in our Apo simulation, quantified by RMSD and structurally depicted in Fig 2. During the initial 900 ns of this simulation, L*β*3*β*4 fluctuates about the “open” crystal structure conformation (small RMSD values, red to light green colors). The transition to the “closed” structure begins to occur at 900 ns (light green to turquoise), seen respectively by a large jump in the loop backbone atom RMSD. The loop begins to sample the “closed” conformation by 1 *µ*s (turquoise to blue), although it is unclear if the transition has completed due to the poor resolution of the “closed” conformation in ZIKV crystal structures.

### Residues of interest for L*β*3*β*4 conformations

Certain residues in the local region of L*β*3*β*4 are hypothesized to have important functions in relation to the loop’s structural states as well as protein-RNA interactions. Such residues are highlighted in Fig 3 and will be discussed in further detail here to present their hypothesized or observed importance.

**Fig 3.**
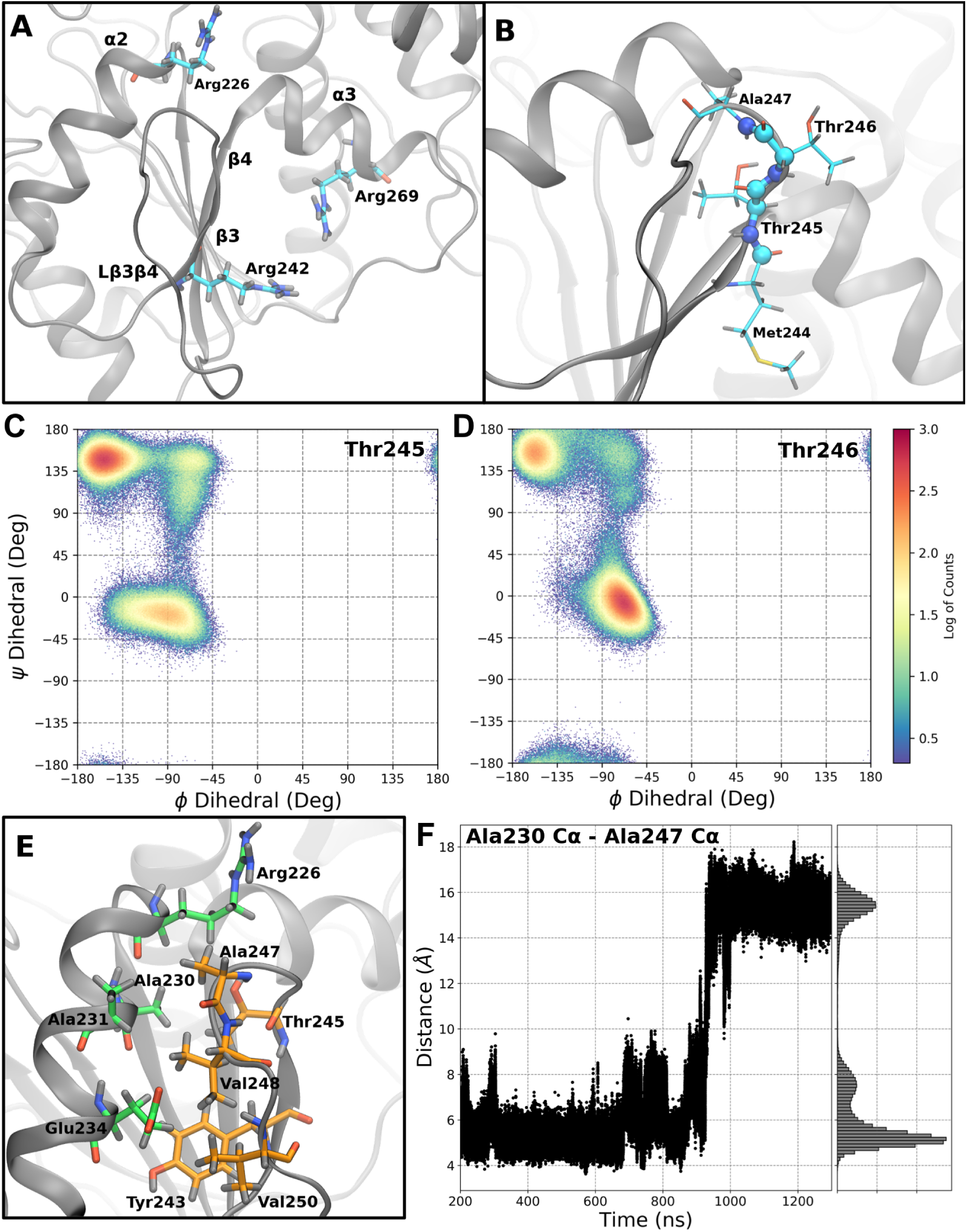
Residues and collective variables that are good descriptors of the “open” to “closed” transition and conformational states. (A) Arginine residues in the RNA-binding cleft, local to L*β*3*β*4. (B) Thr245-Thr246 are residues positioned at the transition between *β*3 and L*β*3*β*4. Ramachandran plots of Thr245 (C) and Thr246 (D) highlight a dihedral switch in these two residues occurring during the loop structural transition. (E) Residues in *α*2 (green) and L*β*3*β*4 (orange) that form a small, stable hydrophobic pocket between the two secondary structures. (F) The time evolution of the distance between Ala230 (*α*2) and Ala247 (L*β*3*β*4) C*α* atoms describes the breakup of the hydrophobic pocket and subsequent loop structural transition.

The three arginine residues highlighted in Fig 3A are ideally positioned within the RNA-binding cleft to function as arginine forks, residues that can strongly coordinate two adjacent phosphate groups of nucleic substrates.^51,52^ Arg226 sits at the top of *α*2, directly interacts with co-crystallized RNA in the 5GJB structure, and is a highly conserved residue of Motif Ia (0.13% sequence variance in flavivirus NS3h sequences).^26^ Numerous crystal structures have resolved the guanidinium functional group of Arg226 in coordination with the RNA phosphate backbone of 3′ terminal nucleotides. Additional arginine residues that are highlighted in panel A, Arg242 (0.67% sequence variance) and Arg269 (1.88% sequence variance), are positioned in *β*3 and *α*3, respectively. These residues have not been observed in crystal structures to coordinate RNA due to the limited resolution of the co-crystallized RNA. Yet, their sequence conservation and ideal positioning within the RNA-binding cleft support our hypothesis that these arginine residues play important roles in binding RNA.

Residues Thr245 and Thr246, shown in Fig 3B, and are positioned at the structural change between *β*3 and L*β*3*β*4. Although less conserved than the residues discussed above, both Thr245 (7.80%) and Thr246 (58.0% sequence variance) are seen to have large structural changes that occur during the “open” to “closed” transition observed in the Apo simulation. Specifically, the loop’s “open” to “closed” conformational change is well described by the simultaneous switching of the backbone dihedrals of these two residues, presented in the Ramachandran plots shown in Fig 3C and D. This observation is supported by these residues’ dihedral values in the flavivirus NS3h crystal structures (see SI). In the “open” conformation of 5JRZ, Thr245 and Thr246 have *ϕ* and *ψ* dihedral values of (−155°, 141°) and (−79°, −1°), respectively. Conversely, in crystal structures with L*β*3*β*4 in the poorly resolved, “closed” conformation (e.g. 5K8L), these residues have *ϕ* and *ψ* dihedral values of (−76°, −12°) and (−141°, 168°), respectively. The Ramachandran plots of Thr245 and Thr246 (Fig 3C and D, respectively) have strong sampling in both dihedral spaces associated with the “open” and “closed” loop conformations. The time evolution of these four dihedrals are shown in Supplementary Figs. 3-6. While the loop shifts from the “open” to the “closed” conformation, large concerted flips in the Thr244 and Thr245 *ψ* dihedrals are observed, followed by correlated fluctuations of these residues’ *ϕ* dihedrals. The dihedral states of these two residues describe the backbone structural conformation of the top of L*β*3*β*4 and so, are hypothesized to be strong decisors of the loop’s conformational state. We hypothesize that these dihedrals will switch during transitions between the “open” and “closed” conformations of the loop for all flavivirus NS3h.

A set of hydrophobic residues in *α*2 and L*β*3*β*4 form a small, hydrophobic pocket when the RNA-binding loop is in the “open” conformation.^27^ This hydrophobic pocket is structurally depicted in Fig 3E, where Ala230 and Ala231 (*α*2) and Ala247, Val248, and Val250 (L*β*3*β*4) are in direct interaction with each other. Additionally, Tyr243 (*β*3) and aliphatic portions of Arg226, Glu234, and Thr245 are neighboring this small hydrophobic region. These stabilizing, hydrophobic interactions are lost in the “closed” conformation of L*β*3*β*4, as quantified by the C*α*-C*α* distance between Ala230 and Ala247 shown in Fig 3F. This distance metric describes the breakup of the depicted hydrophobic pocket and is one of the atomic, pairwise distances with the largest change during the “open” to “closed” transition observed in the Apo simulation.

### Essential dynamics of the L*β*3*β*4 transition

The presented RMSD, dihedral, and distance collective variables clearly delineate the “open” and “closed” conformations of L*β*3*β*4, but there are additional important coordinates necessary to describe the observed transition. To quantify these large-scale motions, the essential dynamics (ED) of L*β*3*β*4 were analyzed. Principal component analysis of the L*β*3*β*4 cartesian coordinates will appropriately quantify the two structural states because the transition between “open” and “closed” conformations represents the largest covariance motions in the coordinate space. Choice of coarse-grained sites for the ED analysis were guided by the multiple sequence analysis results (see SI); residues near the loop region with high sequence conservation were prescribed two sites, describing the backbone and side chain fluctuations independently. Backbone atoms were used to describe the residue-level fluctuations for sequence positions with low conservation or small side chains (e.g. Gly, Ala, and Val). The protocol for determining these atomic coarse-grained sites, detailed in the SI, resulted in 44 atoms that were used to describe the L*β*3*β*4 transition.

The L*β*3*β*4 transition dominates the first eigenvector of the ED analysis, as demonstrated in the scree plot shown in Fig 4A. This eigenvector accounts for 75% of the total variance of the atom selection. The remaining eigenvectors have sufficiently small magnitude eigenvalues and, so, are disregarded for the remainder of this study. The cartesian coordinate data projected onto eigenvector 1 is presented in Fig 4B, where the transition is clearly observed. The “open” conformation is represented by positive projected data values. Additionally, the transition seen at ∼900 ns in Fig 2 is also observed in the projected data. The “closed” conformation is described by large magnitude negative values, with smaller magnitude values representing intermediate loop conformations during the transition.

**Fig 4.**
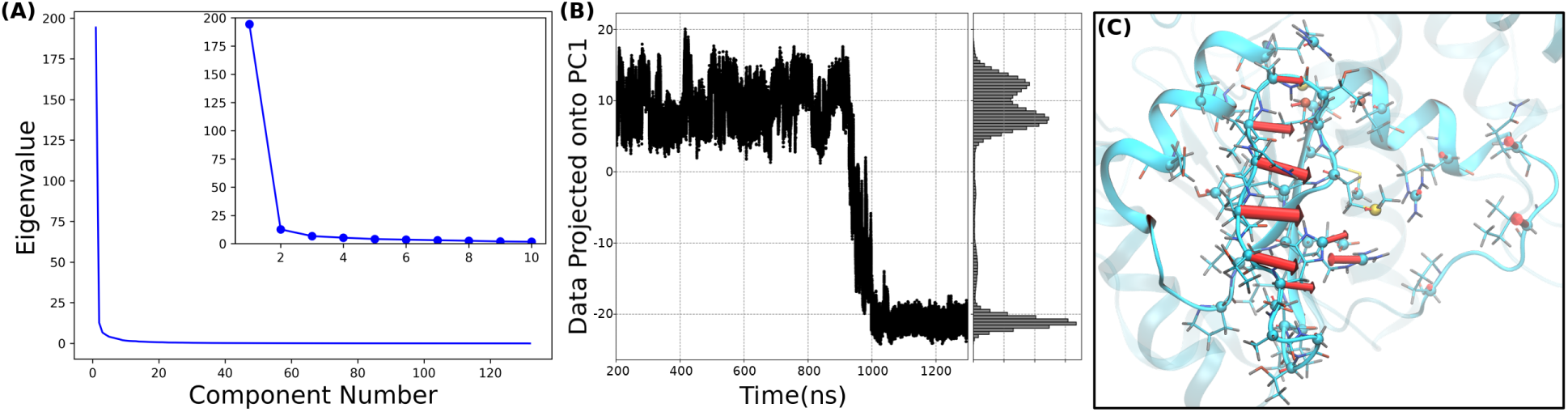
Zika NS3h Apo’s essential dynamics in the L*β*3*β*4 region. (A) The scree plot for the essential dynamics analysis indicates that PC1 is the only major eigenvector to be considered. (B) Projection of the trajectory’s data onto PC1 clearly separates the “open” (positive values) and “closed” (negative values) L*β*3*β*4 conformations. (C) The porcupine plot of the PC1 highlights the correlated fluctuations of the loop residues’ cartesian coordinates during the transition between the two conformational states.

Additionally, the first eigenvector highlights the structural motions that occur during the “open” to “closed” transition. Fig 4C shows the porcupine plot of this eigenvector. The residues with large magnitude vectors in this figure have the largest covariance during the “open” to “closed” conformational change described by PC1. Specifically, C*α* atoms of the loop have the largest magnitude cartesian vectors, indicating strong representation of these residues’ fluctuations in the most dominant eigenvector of the ED analysis. As expected, the direction of these vectors are associated with the “open” to “closed” transition observed in the MD simulation. Interestingly, Arg242 has a large magnitude vector as well pointing in the opposite direction of the loop residues’ large magnitude vectors.

### Free energy landscapes of the RNA-binding loop conformations

We have performed extensive replica exchange umbrella sampling (REUS) simulations to enhance the sampling of the conformational space of L*β*3*β*4 in the Apo, ssRNA, and ssRNA_1−2_ systems. Unfortunately, PC1 cannot be used as the biased collective variable in these REUS simulations due to the non-trivial task of applying a bias on the linear combination of atoms’ coordinates. Instead, the distance between Ala230 (*α*2) and Ala247 (L*β*3*β*4) C*α* atoms was used as proxy for the PC1 reaction coordinate. As previously discussed, this distance represents a large collective variable space and describes the breakup of the hydrophobic interactions between *α*2 and L*β*3*β*4 (see Fig 3). Finally, the dividing surface between the “open” and “closed” conformational states was chosen as the respective collective variable value associated with the highest free energy barrier between the two equilibrium wells. With this dividing surface defined for each system in the Ala230-Ala247 C*α* distance collective variable space, the relative free energy of the two L*β*3*β*4 conformations was approximated by integrating the free energy surface on either side of the divide.

### Relative free energy differences of Apo, ssRNA, and ssRNA_1−2_ systems

The free energy surfaces for the biased CV are shown in Fig 5A, where small (∼4 to 7 Å) and large (∼14 to 18 Å) values correspond to the “open” and “closed” conformations, respectively. In this projection, the Apo substrate state has nearly isoergonic “open” and “closed” conformations (Δ*G*_*O*→*C*_ = −0.22 ± 0.04 kcal mol^−1^), while the two RNA-bound states favor the “open” conformation (Δ*G*_*O*→*C*_ = 0.95 ± 0.02 for ssRNA_1−2_ and 1.97 ± 0.03 kcal mol^−1^ for ssRNA). The small magnitude Δ*G*_*O*→*C*_ for the Apo system corroborates the ensemble of observed RNA-binding loop conformations in crystal structures, where both the “open” and poorly resolved “closed” conformations have been reported numerous times.

**Fig 5.**
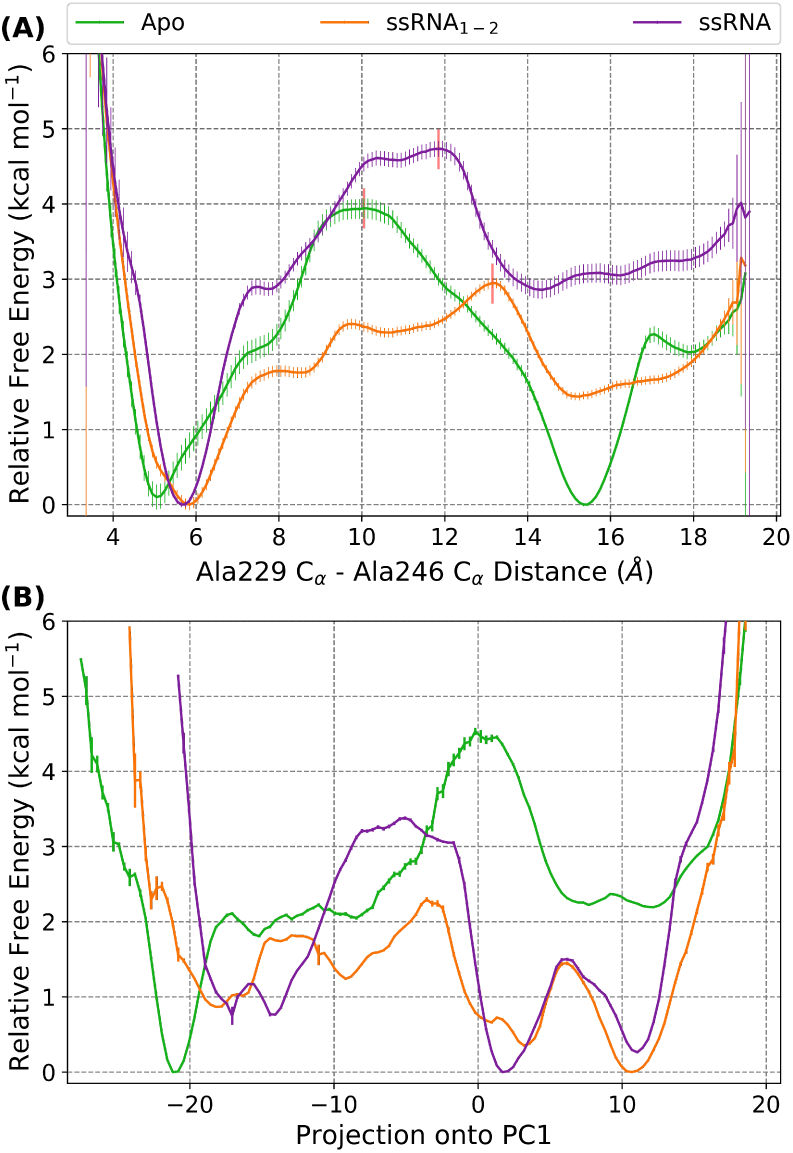
The free energy surfaces for the three systems as projected on the biased CV (panel and PC1 (panel B). (A) Small and large distance values represents the “open” and “closed” conformations, respectively. The transition barrier, shown as a vertical red line, is used to separate the two conformational states of L*β*3*β*4. Error bars were calculated by EMUS, which accounts for the decorrelation time of the collective variable. (B) Positive and negative values correspond to the “closed” and “open” conformations, respectively. Error bars were measured using bootstrapping and so likely under approximate the error in the free energy surfaces.

Both the Δ*G*_*O*→*C*_ and barrier heights shown in Fig 5A highlight the destabilization of the “closed” conformation of L*β*3*β*4 in the presence of RNA. For the ssRNA system, five nucleotides were crystallized in the 5GJB crystal structure. The 3′ end of this RNA oligomer is positioned just at the top of L*β*3*β*4, as seen in Supplementary Fig 2B. The large Δ*G*_*O*→*C*_ and barrier height for the ssRNA system indicate that the close proximity of the 3′ nucleotides locks the RNA-binding loop into the “open” conformation.

Surprisingly, RNA perturbs the loop’s conformational free energy even when the RNA oligomer is *>* 13 Å away from the loop, as is the case for the ssRNA_1−2_ system where three 3′ nucleotides were artificially removed. The remaining, shortened RNA oligomer is positioned between subdomains 2 and 3, between the helical gate *α*-helices. Yet, at this distance, the RNA decreases the free energy barrier between the “open” and “closed” loop conformations relative to both the Apo and ssRNA free energy surfaces. Additionally, this small RNA oligomer shifts the Δ*G*_*O*→*C*_ from favoring the “closed” conformation (as the Apo system does) to favoring the “open” loop.

These results demonstrate that the RNA affects the L*β*3*β*4 free energy surface. In fact, the presence of even minimal RNA at the top of the RNA-binding cleft is seen to perturb the loop’s free energy surface in such a way as to enable the “closed” to “open” transition to occur more readily than in either of the other two substrate states. This may provide insight into the mechanisms of molecular recognition between NS3h and RNA. Specifically, incremental representations of an RNA oligomer bound within NS3h’s RNA-binding cleft are seen to modulate the free energy surface of L*β*3*β*4. Due to its approximately isoergonic Δ*G*_*O*→*C*_ value, the Apo substrate state of NS3h is hypothesized to sample both the “closed” and “open” conformations of the RNA-binding loop; this is strongly supported by the crystal structures of the flavivirus NS3h. The free energy differences of the Apo and ssRNA_1−2_ systems demonstrate that an RNA oligomer positioned between subdomains 2 and 3 (*>* 13 Å away from L*β*3*β*4) induces a population shift away from the “closed” conformation of the loop. This shift is aided by the diminished free energy barrier seen in the ssRNA_1−2_ system’s free energy surface, relative to Apo’s. Further presence of RNA enhances this energetic shift in the free energy surface, as seen in the Δ*G*_*O*→*C*_ for the ssRNA system.

Apo NS3h can sample both the “open” or the “closed” conformations, yet initial binding of RNA drives the free energy surface to favor “open” structures. This results in a population shift away from “closed” structures that is sustained by further binding of the RNA oligomer. Since we do not model the true binding event of RNA into NS3h, we cannot definitively state what mechanism best describes the molecular recognition of RNA by L*β*3*β*4. Yet, the free energy surfaces reported above suggest that the “conformational selection” model of molecular recognition events may be at play for NS3h-RNA complexes, at least in regards to the RNA-binding loop.^53–55^

### Free energy surfaces of the L*β*3*β*4 transition

As stated previously, the biased distance between Ala230 and Ala247 C*α* atoms was used as a proxy to describe the PC1 eigenvector from the Apo system’s ED analysis. Therefore, the REUS simulations were projected onto PC1 to investigate the free energy surface of the loop conformational transition described by this eigenvector. The reweighted free energy surfaces associated with the projection onto PC1 are shown in Fig 5B. For these projected free energy surfaces, less emphasis is placed on the Δ*G*_*O*→*C*_ values due to the larger collective variable space having lower quality sampling for all three systems. In conjunction with the sampling issues in this reweighted space, 1000 iterations of bootstrapping was used to estimate the error in the surfaces; this error is an approximate lower-bound for the true error in these results. Instead, focus is given to the features within these free energy surfaces and their mechanistic interpretations.

#### Apo’s L*β*3*β*4 transition mechanism

The Apo system’s free energy surface in the projected PC1 space (green in Fig 5B) has two major free energy barriers, with the direction of the transition (i.e. “open” to “closed” or the reverse) greatly changing the energetics. From left to right, the first barrier is seen at PC1 ≈ −17 and separates the “closed” conformation from an intermediate “closed” conformation, where the barrier heights of 2.1 or 0.3 kcal mol^−1^ are seen for the “closed” to intermediate and reverse transitions, respectively. Structural representations of these two states are shown in Fig 6A and B, where the loop conformation remains closed, as supported by the dihedral values of Thr245 and Thr246 residues. The largest structural change that separates these two conformations is the Arg242 positioning. In the “closed” conformation, the side chain of this residue sits in a solvent exposed location while, in the intermediate state, the guanidinium group has repositioned into a protein-internal location on the opposite side of the L*β*3*β*4 backbone. This Arg242 repositioning is highlighted in the PC1 porcupine plot (Fig 4C) where the residue’s large magnitude vector is aimed in the opposite direction of the loop’s set of vectors. The Arg242 repositioning represents a large destabilization of the “closed” conformation with a very small barrier in the reverse direction, suggesting that the intermediate “closed” structure is rare and short lived.

**Fig 6.**
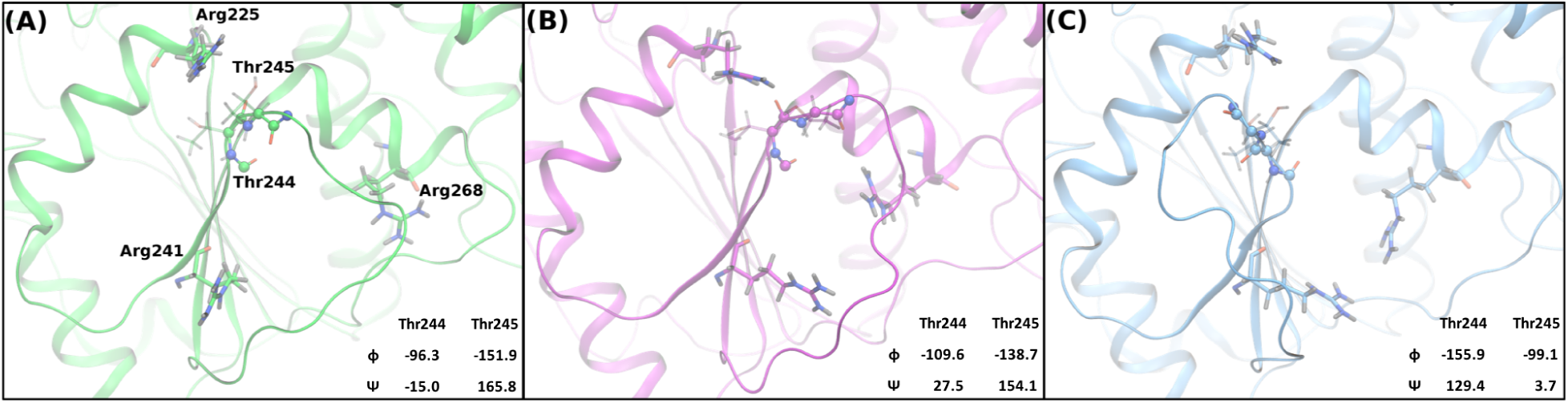
Exemplar structures of the Apo system’s L*β*3*β*4 conformations at local minima in the PC1 projected free energy surface. (A) “Closed” conformation where Arg242 is solvent exposed and Thr245 and Thr246 dihedrals sample the expected “closed” values. (B) The intermediate “closed” conformation where Arg242 has transitioned to the opposite side of L*β*3*β*4, relative to its position in (A). (C) “Open” conformation where the Thr245 and Thr246 dihedrals have flipped into the putative “open” dihedral values and the Arg242 and Arg269 residues sit in the RNA-binding cleft.

The second barrier represents the transition between the intermediate “closed” and “open” conformations, seen at PC1 ≈ 0. An exemplar structure of the “open” conformation is presented in Fig 6C, where the Thr245 and Thr246 dihedrals are seen in the “open” conformation dihedral space. As expected due to the L*β*3*β*4 structure seen in 5JRZ, arginines 226, 242, and 269 are all positioned in the RNA-binding cleft in the “open” conformation. For this transition, the two states are nearly isoergonic with a barrier heights of 2.5 and 2.3 kcal mol^−1^ when transitioning from the intermediate to the “open” structure and the reverse, respectively. With respect to the PC1 porcupine plot, this barrier represents the large structural change of the L*β*3*β*4 residues. The combination of the two barriers seen in the Apo system’s PC1 free energy surface highlights the downhill energetics of the “open” to “closed” transition that was observed in the Apo unbiased simulations.

#### ssRNA systems sample the PC1 transition poorly

Both the ssRNA and ssRNA_1−2_ systems’ PC1-projected free energy surfaces have major differences from the Apo system’s surface in Fig 5B. Important to note are regions on these surfaces with discontinuities or large changes in slope, especially in the transition barriers. The presence of such data indicates non-ergodic sampling at these positions in the PC1 space and suggests that the transition described by PC1 is not viable in the respective substrate state. Even with these caveats, the PC1-projected free energy surfaces of the ssRNA and ssRNA_1−2_ systems are qualitatively similar to the results in the biased CV free energy surfaces. Both systems favor the “open” conformation of L*β*3*β*4 (positive values). The ssRNA_1−2_ oligomer reduces the barrier heights seen in the PC1-projected space relative to those seen in the Apo or ssRNA free energy surfaces.

The ssRNA system’s surface has one large barrier between the two conformational energy wells, approximated as 2.6 and 3.4 kcal mol^−1^ for the “closed” to “open” transition and the reverse, respectively. The transition barrier for this system is the most prominent example of nonergodic sampling and, so, the barrier height is likely under approximated. Furthermore, no intermediate structures between the two wells are seen to be energetically stable and the sampling of negative-valued PC1 space is drastically narrowed. These results suggest that the PC1 eigenvector describes a transition mechanism between “open” and “closed” conformations that is not viable in the ssRNA system. Furthermore, this demonstrates that the energetics of the PC1 transition are strongly affected by the ssRNA oligomer as also observed in the biased CV energetics.

Similarly, the qualitative story observed in the biased CV free energy surface of the ssRNA_1−2_ system is maintained in the PC1-projection surface. The small RNA oligomer stabilizes the “open” conformational state while also minimizing the barrier heights between “closed” and intermediate L*β*3*β*4 structures. The narrowing of PC1 sampling seen for the ssRNA system is not as drastic in the ssRNA_1−2_ system, yet the “closed” conformational well is still displaced to more positive PC1 values. For this system, the structural origins of the RNA oligomer’s effect on the L*β*3*β*4 free energy surface are difficult to identify from the body of simulations reported here. Further study of the numerous RNA-bound structural states of the flavivirus NS3h will need to be performed to fully deconvolute these results.

## Conclusions

The L*β*3*β*4 conformations observed in the flavivirus NS3h crystal structures were investigated using all-atom, explicit solvent MD simulations of the Apo, ssRNA, and ssRNA_1−2_ substrate states. A single observation of the “open” to “closed” conformational transition, seen in the unbiased Apo simulation, was studied to identify collective variables that efficiently quantified the structural changes observed. The backbone dihedrals of the Thr245 and Thr246 residues of L*β*3*β*4 are one such set of collective variables, where crystal structures and MD simulations highlight the large quantitative change in these dihedrals in the “open” and “closed” conformations. Additionally, an essential dynamics analysis of the observed transition produced a single, dominant eigenvector that described 75% of the positional covariance of the RNA-binding loop region. PC1 strongly separated the “open” and “closed” conformations and accounted for the large scale fluctuations of the full loop.

The free energy surfaces of the L*β*3*β*4 structural states were quantified using REUS simulations. Reweighting these free energy results into the PC1 collective variable space also allowed for us to study the energetics of the “open” to “closed” transition originally observed in the unbiased Apo simulation. In either the biased collective variable or PC1 spaces, the quantitative free energy results highlight the RNA-dependence of the L*β*3*β*4 structures. For Apo, a relatively large barrier separates the two structural states that are nearly isoergonic in the Ala230-Ala247 C*α* distance space. In the projected space, the transition from “open” to “closed” is much more energetically favorable than the reverse process. In this transition, two energetic steps are observed to occur: the backbone of the RNA-binding loop moves into the “open” conformation, as described by the Thr245 and Thr246 dihedrals, followed by the side chain of Arg242 fluctuating to the opposite side of the loop to a position in the RNA-binding cleft.

For the ssRNA and ssRNA_1−2_ systems, the “open” loop conformation is energetically favored in both free energy surfaces. Direct interactions between RNA and loop residues lead to an even larger barrier seen for the ssRNA system’s biased collective variable free energy surface. Additionally, the transition described by PC1 is not viable in the ssRNA system as indicated by the nonergodic regions in the respective surface. Interestingly, the ssRNA_1−2_ system has lower free energy barriers in both the biased collective variable and PC1 free energy surfaces. These results suggest that even a small RNA oligomer, bound in the RNA-binding cleft between subdomains 2 and 3, increases the rate at which L*β*3*β*4 transitions between the “closed” conformation (favored by Apo) and the “open” conformation. These results also suggest a possible conformational selection mechanism for the RNA-dependent L*β*3*β*4 structural states, where RNA positioned between subdomains 2 and 3 causes an energetic shift in the loop’s structures to favor the RNA-bound, “open” conformation.

In light of our previous study of the dengue NS3h NTPase cycle, the L*β*3*β*4’s RNA-dependent free energy surface provides interesting insight into the hypothesis that RNA-induced enhancement of the NTP hydrolysis reaction originates, to some degree, from the “open” L*β*3*β*4 conformation. We previously reported that the number of water molecules within the hydrolysis active site is decreased as are these waters’ translational and rotational motions when RNA is present.^27^ Additionally, the positioning of water molecules within the active site was also observed to be affected by RNA. These effects are proposed to originate from RNA-induced structural changes in the hydrolysis active site, an example of which is a structural shift in *α*2 brought on by the L*β*3*β*4 interactions while in the “open” conformation. Therefore, our free energy results suggest that initial binding states of the NS3h:RNA complex (modeled by ssRNA_1−2_) enhance the L*β*3*β*4 transition from “closed” to “open”, leading to structural shifts in *α*2 that prime the NTPase active site for the hydrolysis reaction. Continued binding of RNA (modeled by ssRNA) locks in the *α*2-L*β*3*β*4 interactions even further.

The structural origins of the ssRNA_1−2_ oligomer’s effect on L*β*3*β*4 have not been identified, yet our results highlight the RNA-binding loop as a potential target for small-molecule inhibition or mutagenesis studies. We hypothesize that destabilization or prevention of the “open” L*β*3*β*4 structure, while in the Apo or ssRNA_1−2_ substrate states, would diminish the RNA-induced enhancement of the NTPase activity as well as lead to weakened protein-RNA interactions. The highly conserved Arg226, Arg242, and Arg269 are all hypothesized to function as arginine fork residues that strongly coordinate the phosphate groups of 3′ RNA nucleotides. Arg242 is also observed to play an important role in the “open” to “closed” transition described by PC1; stabilization of the solvent exposed conformation of Arg242 (seen in Fig 6) is proposed as a potential method of inhibiting the transition. Therefore, mutations or small molecules that perturb the wild-type behavior of L*β*3*β*4 would be detectable via NTPase, RNA binding, and helicase activity assays. Additionally, further computational studies of the NS3h:RNA structural states could provide insights into the short-range, residue-residue interactions through which the observed allosteric effect is propagated.

## Supplementary Information

**S1 File. Supplementary information file.** The file includes the weblogo and structural representations of the flavivirus NS3h MSA results and structural alignments. Additionally, time evolution plots of the Thr245 and Thr246 backbone dihedrals are presented. (PDF)

## Acknowledgments

RBD would like to acknowledge Dr. Peter Lake for insightful discussions.

## References

1. Bhatt, S.; Gething, P. W.; Brady, O. J.; Messina, J. P.; Farlow, A. W.; Moyes, C. L.; Drake, J. M.; Brownstein, J. S.; Hoen, A. G.; Sankoh, O.; Myers, M. F.; George, D. B.; Jaenisch, T.; Wint, G. R. W.; Simmons, C. P.; Scott, T. W.; Farrar, J. J.; Hay, S. I. The global distribution and burden of dengue. Nature 2013, 496, 504–507.

2. Messina, J. P.; Kraemer, M. U.; Brady, O. J.; Pigott, D. M.; Shearer, F. M.; Weiss, D. J.; Golding, N.; Ruktanonchai, C. W.; Gething, P. W.; Cohn, E.; Brownstein, J. S.; Khan, K.; Tatem, A. J.; Jaenisch, T.; Murray, C. J.; Marinho, F.; Scott, T. W.; Hay, S. I. Mapping global environmental suitability for Zika virus. eLife 2016, 5, e15272.

3. Musso, D.; Gubler, D. J. Zika Virus. Clin. Microbiol. Rev. 2016, 29, 487–524.

4. Ekins, S.; Perryman, A. L.; Horta Andrade, C. OpenZika: An IBM World Community Grid Project to Accelerate Zika Virus Drug Discovery. PLoS Neg. Trop. Dis. 2016, 10, e0005023.

5. Ekins, S.; Mietchen, D.; Coffee, M.; Stratton, T. P.; Freundlich, J. S.; Freitas-Junior, L.; Muratov, E.; Siqueira-Neto, J.; Williams, A. J.; Andrade, C. Open drug discovery for the Zika virus. F1000Res 2016, 5, 150.

6. Gupta, A. K.; Kaur, K.; Rajput, A.; Dhanda, S. K.; Sehgal, M.; Khan, M. S.; Monga, I.; Dar, S. A.; Singh, S.; Nagpal, G.; Usmani, S. S.; Thakur, A.; Kaur, G.; Sharma, S.; Bhardwaj, A.; Qureshi, A.; Raghava, G. P. S.; Kumar, M. ZikaVR: An Integrated Zika Virus Resource for Genomics, Proteomics, Phylogenetic and Therapeutic Analysis. Sci. Rep. 2016, 6, 32713.

7. Cox, B. D.; Stanton, R. A.; Schinazi, R. F. Predicting Zika virus structural biology: Challenges and opportunities for intervention. Antivir. Chem. Chemother. 2015, 24, 118–126.

8. Leung, D.; Schroder, K.; White, H.; Fang, N. X.; Stoermer, M. J.; Abbenante, G.; Martin, J. L.; Young, P. R.; Fairlie, D. P. Activity of Recombinant Dengue 2 Virus NS3 Protease in the Presence of a Truncated NS2B Co-factor, Small Peptide Substrates, and Inhibitors. J. Biol. Chem. 2001, 276, 45762–45771.

9. Borowski, P.; Niebuhr, A.; Schmitz, H.; Hosmane, R. S.; Bretner, M.; Siwecka, M. A.; Kulikowski, T. NTPase/helicase of Flaviviridae: inhibitors and inhibition of the enzyme. Acta Biochim. Pol. 2002, 49, 597–614.

10. Raney, K. D.; Sharma, S. D.; Moustafa, I. M.; Cameron, C. E. Hepatitis C virus non-structural protein 3 (HCV NS3): a multifunctional antiviral target. J. Biol. Chem. 2010, 285, 22725–31.

11. Byrd, C. M.; Grosenbach, D. W.; Berhanu, A.; Dai, D.; Jones, K. F.; Cardwell, K. B.; Schneider, C.; Yang, G.; Tyavanagimatt, S.; Harver, C.; Wineinger, K. A.; Page, J.; Stavale, E.; Stone, M. A.; Fuller, K. P.; Lovejoy, C.; Leeds, J. M.; Hruby, D. E.; Jordan, R. Novel benzoxazole inhibitor of dengue virus replication that targets the NS3 helicase. Antimicrob. Agents Chemother. 2013, 57, 1902–1912.

12. Basavannacharya, C.; Vasudevan, S. G. Suramin inhibits helicase activity of NS3 protein of dengue virus in a fluorescence-based high throughput assay format. Biochem. Biophys. Res. Commun. 2014, 453, 539–44.

13. Ndjomou, J.; Corby, M. J.; Sweeney, N. L.; Hanson, A. M.; Aydin, C.; Ali, A.; Schiffer, C. A.; Li, K.; Frankowski, K. J.; Schoenen, F. J.; Frick, D. N. Simultaneously Targeting the NS3 Protease and Helicase Activities for More Effective Hepatitis C Virus Therapy. ACS Chem. Biol. 2015, 10, 1887–1896.

14. Sweeney, N. L.; Hanson, A. M.; Mukherjee, S.; Ndjomou, J.; Geiss, B. J.; Steel, J. J.; Frankowski, K. J.; Li, K.; Schoenen, F. J.; Frick, D. N. Benzothiazole and Pyrrolone Flavivirus Inhibitors Targeting the Viral Helicase. ACS Infect. Dis. 2015, 1, 140–148.

15. Lee, H.; Ren, J.; Nocadello, S.; Rice, A. J.; Ojeda, I.; Light, S.; Minasov, G.; Vargas, J.; Nagarathnam, D.; Anderson, W. F.; Johnson, M. E. Identification of novel small molecule inhibitors against NS2B/NS3 serine protease from Zika virus. Antiviral Res. 2017, 139, 49–58.

16. Munawar, A.; Beelen, S.; Munawar, A.; Lescrinier, E.; Strelkov, S.; Munawar, A.; Beelen, S.; Munawar, A.; Lescrinier, E.; Strelkov, S. V. Discovery of Novel Druggable Sites on Zika Virus NS3 Helicase Using X-ray Crystallography-Based Fragment Screening. Int. J. Mol. Sci. 2018, 19, 3664.

17. Kinney, R. M.; Butrapet, S.; Chang, G. J. J.; Tsuchiya, K. R.; Roehrig, J. T.; Bhamarapravati, N.; Gubler, D. J. Construction of infectious cDNA clones for dengue 2 virus: Strain 16681 and its attenuated vaccine derivative, strain PDK-53. Virology 1997, 230, 300–308.

18. Osorio, J. E.; Huang, C. Y. H.; Kinney, R. M.; Stinchcomb, D. T. Development of DENVax: A chimeric dengue-2 PDK-53-based tetravalent vaccine for protection against dengue fever. Vaccine 2011, 29, 7251–7260.

19. Wengler, G.; Wengler, G. The carboxy-terminal part of the NS 3 protein of the West Nile flavivirus can be isolated as a soluble protein after proteolytic cleavage and represents an RNA-stimulated NTPase. Virology 1991, 184, 707–715.

20. Warrener, P.; Tamura, J. K.; Collett, M. S. RNA-stimulated NTPase activity associated with yellow fever virus NS3 protein expressed in bacteria. J. Virol. 1993, 67, 989–996.

21. Kuo, M. D.; Chin, C.; Hsu, S. L.; Shiao, J. Y.; Wang, T. M.; Lin, J. H. Characterization of the NTPase activity of Japanese encephalitis virus NS3 protein. J. Gen. Virol. 1996, 77, 2077–2084.

22. Utama, A.; Shimizu, H.; Morikawa, S.; Hasebe, F.; Morita, K.; Igarashi, A.; Hatsu, M.; Takamizawa, K.; Miyamura, T. Identification and characterization of the RNA helicase activity of Japanese encephalitis virus NS3 protein. FEBS Lett. 2000, 465, 74–78.

23. Borowski, P.; Niebuhr, A.; Mueller, O.; Bretner, M.; Felczak, K.; Kulikowski, T.; Schmitz, H. Purification and characterization of West Nile virus nucleoside triphosphatase (NTPase)/helicase: evidence for dissociation of the NTPase and helicase activities of the enzyme. J. Virol. 2001, 75, 3220–3229.

24. Wang, C.-C.; Huang, Z.-S.; Chiang, P.-L.; Chen, C.-T.; Wu, H.-N. Analysis of the nucleoside triphosphatase, RNA triphosphatase, and unwinding activities of the helicase domain of dengue virus NS3 protein. FEBS Lett. 2009, 583, 691–696.

25. Klema, V. J.; Padmanabhan, R.; Choi, K. H. Flaviviral Replication Complex: Coordination between RNA Synthesis and 51-RNA Capping. Viruses 2015, 7, 4640–4656.

26. Fairman-Williams, M. E.; Guenther, U.-P.; Jankowsky, E. SF1 and SF2 helicases: family matters. Curr. Opin. Struct. Biol. 2010, 20, 313–324.

27. Davidson, R. B.; Hendrix, J.; Geiss, B. J.; McCullagh, M. Allostery in the dengue virus NS3 helicase: Insights into the NTPase cycle from molecular simulations. PLoS Comput. Biol. 2018, 14, e1006103–28.

28. Swarbrick, C. M.; Basavannacharya, C.; Chan, K. W.; Chan, S. A.; Singh, D.; Wei, N.; Phoo, W. W.; Luo, D.; Lescar, J.; Vasudevan, S. G. NS3 helicase from dengue virus specifically recognizes viral RNA sequence to ensure optimal replication. Nucleic Acids Res. 2017, 45, 12904–12920.

29. Jain, R.; Coloma, J.; Garcia-Sastre, A.; Aggarwal, A. K. Structure of the NS3 helicase from Zika virus. Nat. Struct. Mol. Biol. 2016, 23, 752–754.

30. Cao, X.; Li, Y.; Jin, X.; Li, Y.; Guo, F.; Jin, T. Molecular mechanism of divalent-metal-induced activation of NS3 helicase and insights into Zika virus inhibitor design. Nucleic Acids Res. 2016, 44, 10505–10514.

31. Mottin, M.; Braga, R. C.; da Silva, R. A.; da Silva, J. H.; Perryman, A. L.; Ekins, S.; Andrade, C. H. Molecular dynamics simulations of Zika virus NS3 helicase: Insights into RNA binding site activity. Biochem. Biophys. Res. Commun. 2017, 492, 643–651.

32. Du Pont, K. E.; Davidson, R. B.; McCullagh, M.; Geiss, B. J. Motif V regulates energy transduction between the flavivirus NS3 ATPase and RNA-binding cleft. J. Biol. Chem. 2019, jbc.RA119.011922.

33. Gu, M.; Rice, C. M. The Spring *α*-Helix Coordinates Multiple Modes of HCV (Hepatitis C Virus) NS3 Helicase Action. J. Biol. Chem. 2016, 291, 14499–509.

34. Tian, H.; Ji, X.; Yang, X.; Zhang, Z.; Lu, Z.; Yang, K.; Chen, C.; Zhao, Q.; Chi, H.; Mu, Z.; Xie, W.; Wang, Z.; Lou, H.; Yang, H.; Rao, Z. Structural basis of Zika virus helicase in recognizing its substrates. Protein & Cell 2016, 7, 562–570.

35. Luo, D.; Xu, T.; Watson, R. P.; Scherer-Becker, D.; Sampath, A.; Jahnke, W.; Yeong, S. S.; Wang, C. H.; Lim, S. P.; Strongin, A.; Vasudevan, S. G.; Lescar, J. Insights into RNA unwinding and ATP hydrolysis by the flavivirus NS3 protein. EMBO J. 2008, 27, 3209–3219.

36. Case, D. A.; Betz, R.; Cerutti, D.; Cheatham, T. I.; Darden, T.; Duke, R.; Giese, T.; Gohlke, H.; Goetz, A.; Homeyer, N.; Izadi, S.; Janowski, P.; Kaus, J.; Kovalenko, A.; Lee, T.; LeGrand, S.; Li, P.; Lin, C.; Luchko, T.; Luo, R.; Madej, B.; Mermelstein, D.; Merz, K.; Monard, G.; Nguyen, H.; Nguyen, H.; Omelyan, I.; Onufriev, A.; Roe, D.; Roitberg, A.; Sagui, C.; Simmerling, C.; Botello-Smith, W.; Swails, J.; Walker, R.; Wang, J.; Wolf, R.; Wu, X.; Xiao, L.; Kollman, P. AMBER16. 2016.

37. Case, D. A.; Ben-Shalom, I.; Brozell, S.; Cerutti, D.; Cheatham, T. I.; Cruzeiro, V.; Darden, T.; Duke, R.; Ghoreishi, D.; Gilson, M.; Gohlke, H.; Goetz, A.; Greene, D.; Harris, R.; Homeyer, N.; Izadi, S.; Kovalenko, A.; Kurtzman, T.; Lee, T.; LeGrand, S.; Li, P.; Lin, C.; Liu, J.; Luchko, T.; Luo, R.; Mermelstein, D.; Merz, K.; Miao, Y.; Monard, G.; Nguyen, C.; Nguyen, H.; Omelyan, I.; Onufriev, A.; Pan, F.; Qi, R.; Roe, D.; Roitberg, A.; Sagui, C.; Schott-Verdugo, S.; Shen, J.; Simmerling, C.; Smith, J.; Salomon-Ferrer, R.; Swails, J.; Walker, R.; Wang, J.; Wei, H.; Wolf, R.; Wu, X.; Xiao, L.; York, D.; Kollman, P. AMBER18. 2018.

38. Maier, J. A.; Martinez, C.; Kasavajhala, K.; Wickstrom, L.; Hauser, K. E.; Simmerling, C. ff14SB: Improving the Accuracy of Protein Side Chain and Backbone Parameters from ff99SB. J. Chem. Theory Comput. 2015, 11, 3696–3713.

39. Banáš, P.; Hollas, D.; Zgarbová, M.; Jurečka, P.; Orozco, M.; Cheatham, T. E.; Šponer, J.; Otyepka, M. Performance of Molecular Mechanics Force Fields for RNA Simulations: Stability of UUCG and GNRA Hairpins. J. Chem. Theory Comput. 2010, 6, 3836–3849.

40. Zgarbová, M.; Otyepka, M.; Šponer, J.; Mládek, A.; Banáš, P.; Cheatham, T. E.; Jurečka, P. Refinement of the Cornell et al. Nucleic Acids Force Field Based on Reference Quantum Chemical Calculations of Glycosidic Torsion Profiles. J. Chem. Theory Comput. 2011, 7, 2886–2902.

41. Ryckaert, J.-P.; Ciccotti, G.; Berendsen, H. J. Numerical integration of the cartesian effiuations of motion of a system with constraints: molecular dynamics of n-alkanes. J. Comp. Phys. 1977, 23, 327–341.

42. Cheatham, T. E. I.; Miller, J. L.; Fox, T.; Darden, T. A.; Kollman, P. A. Molecular Dynamics Simulations on Solvated Biomolecular Systems: The Particle Mesh Ewald Method Leads to Stable Trajectories of DNA, RNA, and Proteins. J. Am. Chem. Soc. 1995, 117, 4193–4194.

43. Isralewitz, B.; Gao, M.; Schulten, K. Steered molecular dynamics and mechanical functions of proteins. 2001, 11, 224–230.

44. Fukunishi, H.; Watanabe, O.; Takada, S. On the Hamiltonian replica exchange method for efficient sampling of biomolecular systems: Application to protein structure prediction. J. Chem. Phys. 2002, 116, 9058–9067.

45. Thiede, E. H.; Van Koten, B.; Weare, J.; Dinner, A. R. Eigenvector method for umbrella sampling enables error analysis. J. Chem. Phys. 2016, 145, 084115.

46. Michaud-Agrawal, N.; Denning, E. J.; Woolf, T. B.; Beckstein, O. MDAnalysis: A toolkit for the analysis of molecular dynamics simulations. J. Comput. Chem. 2011, 32, 2319–2327.

47. Hunter, J. D. Matplotlib: A 2D graphics environment. Comput. Sci. Eng. 2007, 9, 90–95.

48. Humphrey, W.; Dalke, A.; Schulten, K. VMD: visual molecular dynamics. J. Mol. Graph. 1996, 14, 33–38.

49. Frishman, D.; Argos, P. Knowledge-based protein secondary structure assignment. Proteins 1995, 23, 566–579.

50. Stone, J. An Efficient Library for Parallel Ray Tracing and Animation. M.Sc. thesis, Computer Science Department, University of Missouri-Rolla, 1998.

51. Calnan, B. J.; Tidor, B.; Biancalana, S.; Hudson, D.; Frankel, A. D. Arginine-mediated RNA recognition: The arginine fork. Science 1991, 252, 1167–1171.

52. Tao, J.; Frankel, A. D. et al. Specific binding of arginine to TAR RNA. Proc. Natl. Acad. Sci. U.S.A. 1992, 89, 2723–2726.

53. Tsai, C. J.; Ma, B.; Nussinov, R. Folding and binding cascades: Shifts in energy landscapes. Proc. Natl. Acad. Sci. U.S.A. 1999, 96, 9970–9972.

54. Boehr, D. D.; Nussinov, R.; Wright, P. E. The role of dynamic conformational ensembles in biomolecular recognition. Nat. Chem. Biol. 2009, 5, 789–796.

55. Kumar, S.; Ma, B.; Tsai, C.-J.; Sinha, N.; Nussinov, R. Folding and binding cascades: Dynamic landscapes and population shifts. Protein Sci. 2008, 9, 10–19.

