## Supporting Information for "RNA-dependent structures of the RNA-binding loop in the flavivirus NS3 helicase"

### Supporting Information for:
### RNA-dependent structures of the RNA-binding loop in the flavivirus NS3 helicase

Russell B. Davidson, Josie Hendrix, Brian J. Geiss, Martin McCullagh

#### Analyses

##### Multiple Sequence Alignment

The NIAID Virus Pathogen Database and Analysis Resource (ViPR) [1] (accessed on May 30th, 2018 through the web site at http://www.viprbrc.org/) was used to collect the available *Flaviviridae* NS3 sequences, totaling 51,816 independent sequences from all four genuses of the virus family. The Biopython module [2] (version 1.74) was used to preprocess this population of NS3 sequences. Specifically, this preprocessing analysis removed uninteresting sequences (not an NS3h sequence, too short to be full NS3h, or poorly resolved). Subsequently, the CD-HIT software package was used to perform a three-step clustering analysis. [3,4] The first clustering step used a clustering threshold of 1.00, which was used to aggregate non-unique sequences into a single, representative sequence. This was done to normalize the weight that each unique sequence has in the population of sequences. The second step in the sequence clustering protocol used a clustering threshold of 0.4, which successfully clustered sequences by the genera of the *Flaviviridae* family (flavivirus, hepacivirus, pestivirus, and pegivirus). Finally, the third step of clustering was performed on the flavivirus sequences with a threshold of 0.8. This last step was used to remove poorly sampled clusters of sequences, which were either poor quality sequences or associated with flaviviruses (e.g. Apoi virus) that had few representative sequences. After the preprocessing analysis and clustering protocol, 1,488 flaviviral sequences remained and were subsequently analyzed in a multiple sequence alignment (MSA).

The MAFFT software [5] was used to perform three independent MSA analyses: (1) a progressive alignment with iterative refinement, (2) local alignment with iterative refinement methods using WSP and consistency scores, and (3) global alignment with iterative refinement methods using WSP and consistency scores. Output from these separate MSA analyses were then analyzed in the trimAl software [6] to quantitatively chose the highest quality MSA as well as remove sequence positions with gaps in at least 20% of the analyzed sequences. Finally, post-analysis of the MSA results measured the sequence position variance away from the consensus sequence. Specifically, the position frequency matrix was calculated from the ensemble of aligned sequences. The variance away from the most-probable residue at each sequence position was calculated from this data; these values are reported in the paper for specific residues of interest. The MSA results are provided in the zipped Supplementary Information (SI) directory. Additionally, Fig 1 depicts the sequence logo of the MSA results, using dengue NS3h residue numbering. [7,8] The exact pre-processing, clustering, MSA, and post-processing protocols are available on Github (`https://github.com/mccullaghlab/ZIKV-Lb3b4/bioinformatics`).

Fig 2A highlights the sequence variance of sequence positions in the local region of L$\beta3\beta4$. Sequence positions with high conservation (< 1% variance) across all flavivirus sequences are colored blue, while positions with larger variance away from the consensus are colored from white to red, representing increasing variance. This structural representation of the MSA results highlight the poor conservation of the loop residues while sequence positions in $\alpha2$, $\beta3$, $\beta4$, and $\alpha3$ secondary structures are more strongly conserved.

##### Crystal Structure Alignment

In conjunction with the MSA analysis of flavivirus NS3h sequences, a structural alignment of all available crystal structures of the flavivirus NS3h was performed to highlight the structural heterogeneity of the RNA-binding loop and surrounding areas. In total, 56 monomer structures of flavivirus NS3h are available on the Protein Data Bank (PDB). The dengue NS3h:ssRNA:ATP crystal structure (2JLV) was used as the alignment reference structure. The alignment landmark reported in Davidson *et al.* [9] was used. Specifically, the C$\alpha$ atoms of the core $\beta$-sheets of subdomains 1 and 2 were used for alignment due to the strongly maintained tertiary structure of these subdomains. The aligned structures and VMD visualization states are provided in the zipped SI directory.

Panel B of Fig 2 depicts a representative body of crystal conformations of ZIKV NS3h, focusing on the L$\beta3\beta4$ structural region. Generally, the "closed" conformation of the loop is poorly resolved except for the initial residues of the loop. Conversely, structures with the loop in the "open" conformation are well resolved and have similar structures to the RNA-bound (5GJB)

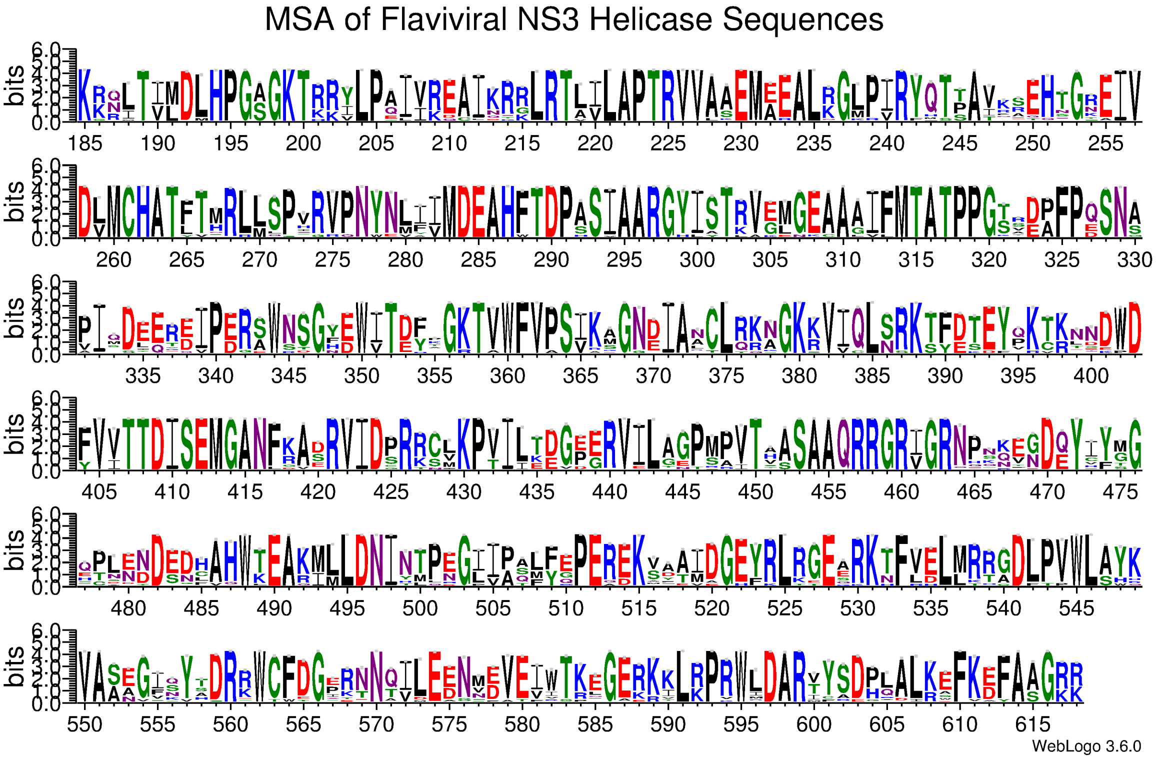


**Fig 1.** Sequence logo[7,8] of the flaviviral NS3h multiple sequence alignment results, using residue numbering from the DENV NS3h 2JLV structure. An equiprobable background composition of amino acid usage was assumed. Amino acids are colored based on their side chain chemistry: polar residues (green), neutral (purple), basic (blue), acidic (red), and hydrophobic (black). The relative height of each residue letter describes the relative frequency of observing the respective residue at that sequence position. The overall height of the column describes how conserved the sequence position is.

crystal. Panels C and D of this figure highlight the arginine fork residues found in the L$\beta3\beta4$ local region of the RNA-binding cleft as well as the Thr245 and Thr246 residues, which are seen in structural analyses to be strong decisors of the two conformational states of the loop.

#### Essential Dynamics of the L$\beta3\beta4$ Structure

The largest covariance motions of the RNA-binding loop were analyzed in the unbiased Apo simulation. Specifically, residues in $\alpha2$, $\beta3$, L$\beta3\beta4$, $\beta4$, and $\alpha3$ were of interest when quantifying the "open" to "closed" conformational change observed in the unbiased simulation. Of these secondary structures, residues that face away from the RNA-binding cleft were not considered. The MSA results guided the choice of coarse-graining used in this essential dynamics (ED) analysis. If a sequence position is strongly conserved (low sequence position variance), then the residue fluctuations are described with a two site coarse-graining: the C$\alpha$ atom used to describe backbone fluctuations and a side chain atom to describe the side chain fluctuations. An exception to this are residues with small side chains (e.g. Gly, Ala, and Val) where the fluctuation of the side chain is highly correlated with the backbone fluctuations. Residues with low conservation are coarse-grained as a single site at the C$\alpha$ atom. This bioinformatics-guided coarse-graining resulted in 44 atoms, incorporating all five secondary structures in the region of interest. A principal component analysis (PCA) of these atoms' cartesian coordinates result in eigenvectors describing the essential dynamics of the L$\beta3\beta4$ conformational transition observed in the Apo simulation.

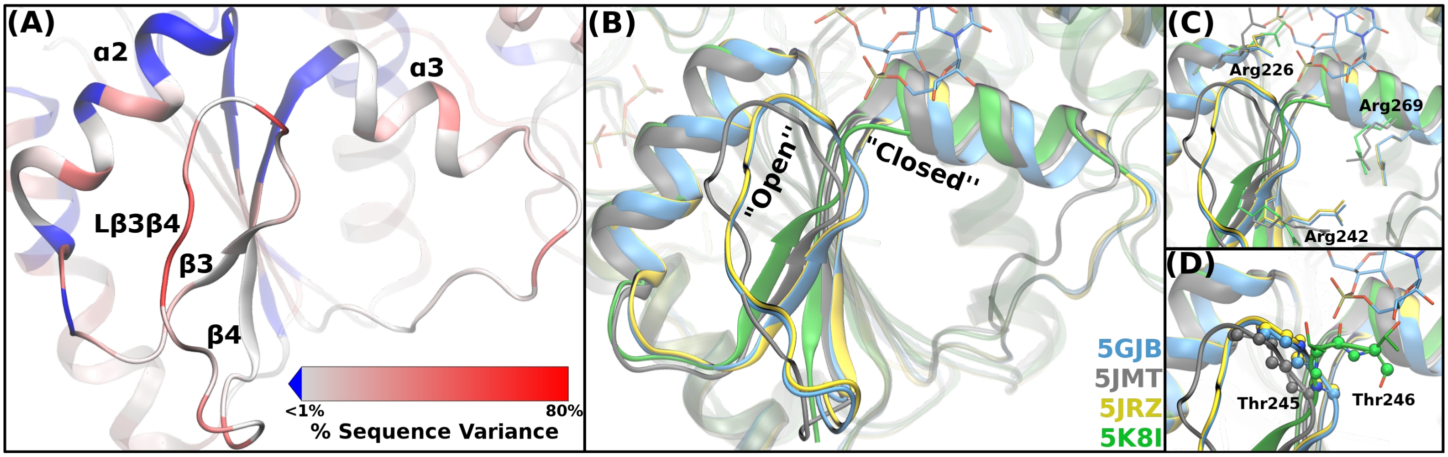


**Fig 2.** Alignment of flaviviral NS3h sequences and crystal structures. (A) Structural representation of the MSA results. Sequence positions are colored based on percent variance away from the consensus sequence. Highly conserved positions are colored blue, while less conserved residues are colored from white to red with increasing variance. (B) Crystal structure alignment of a subset of ZIKV NS3h, focusing on the L$\beta3\beta4$ region of subdomain 1. 5K8I is one of the few crystal structures with the L$\beta3\beta4$ structure in a "closed"-like position, albeit largely unresolved. (B) The strongly conserved Arg226, Arg242, and Arg269 residues hypothesized to function as arginine forks. (C) The large L$\beta3\beta4$ structural change can be seen in the large dihedral shifts of the Thr245-Thr246 residue pair.

### References

1. Pickett, B. E.; Sadat, E. L.; Zhang, Y.; Noronha, J. M.; Squires, R. B.; Hunt, V.; Liu, M.; Kumar, S.; Zaremba, S.; Gu, Z.; Zhou, L.; Larson, C. N.; Dietrich, J.; Klem, E. B.; Scheuermann, R. H. ViPR: an open bioinformatics database and analysis resource for virology research. *Nucleic Acids Res.* **2012**, *40*, D593–D598.

2. Cock, P. J.; Antao, T.; Chang, J. T.; Chapman, B. A.; Cox, C. J.; Dalke, A.; Friedberg, I.; Hamelryck, T.; Kauff, F.; Wilczynski, B.; De Hoon, M. J. Biopython: Freely available Python tools for computational molecular biology and bioinformatics. *Bioinformatics* **2009**, *25*, 1422–1423.

3. Fu, L.; Niu, B.; Zhu, Z.; Wu, S.; Li, W. CD-HIT: Accelerated for clustering the next-generation sequencing data. *Bioinformatics* **2012**, *28*, 3150–3152.

4. Li, W.; Godzik, A. Cd-hit: A fast program for clustering and comparing large sets of protein or nucleotide sequences. *Bioinformatics* **2006**, *22*, 1658–1659.

5. Katoh, K.; Misawa, K.; Kuma, K.; Miyata, T. MAFFT: a novel method for rapid multiple sequence alignment based on fast Fourier transform. *Nucleic Acids Res.* **2002**, *30*, 3059–3066.

6. Capella-Gutiérrez, S.; Silla-Martínez, J. M.; Gabaldón, T. trimAl: A tool for automated alignment trimming in large-scale phylogenetic analyses. *Bioinformatics* **2009**, *25*, 1972–1973.

7. Crooks, G. E.; Hon, G.; Chandonia, J.-M.; Brenner, S. E. WebLogo: A Sequence Logo Generator. *Genome Res.* **2004**, *14*, 1188–1190.

8. Schneider, T. D.; Stephens, R. M. Sequence logos: A new way to display consensus sequences. *Nucleic Acids Res.* **1990**, *18*, 6097–6100.

9. Davidson, R. B.; Hendrix, J.; Geiss, B. J.; McCullagh, M. Allostery in the dengue virus NS3 helicase: Insights into the NTPase cycle from molecular simulations. *PLoS Comput. Biol.* **2018**, *14*, e1006103–28.

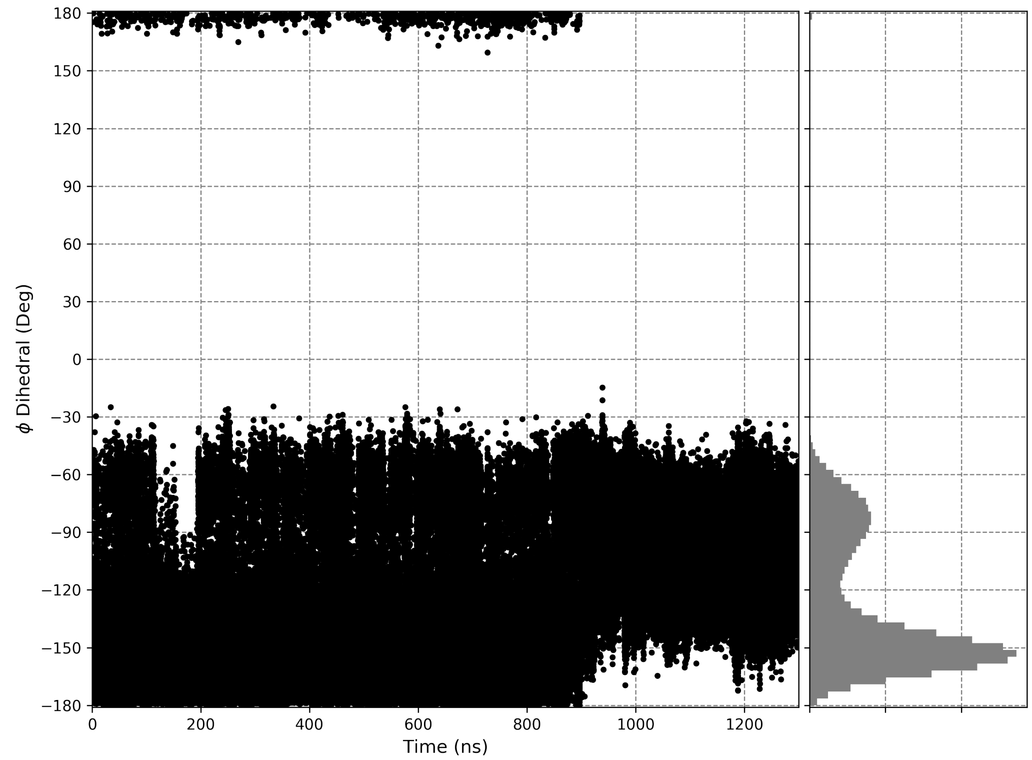


**Fig 3.** $\phi$ dihedral of Thr245 in the unbiased Apo simulation. The "open" to "closed" conformational change in L$\beta$3$\beta$4 occurs at approximately 900 ns.

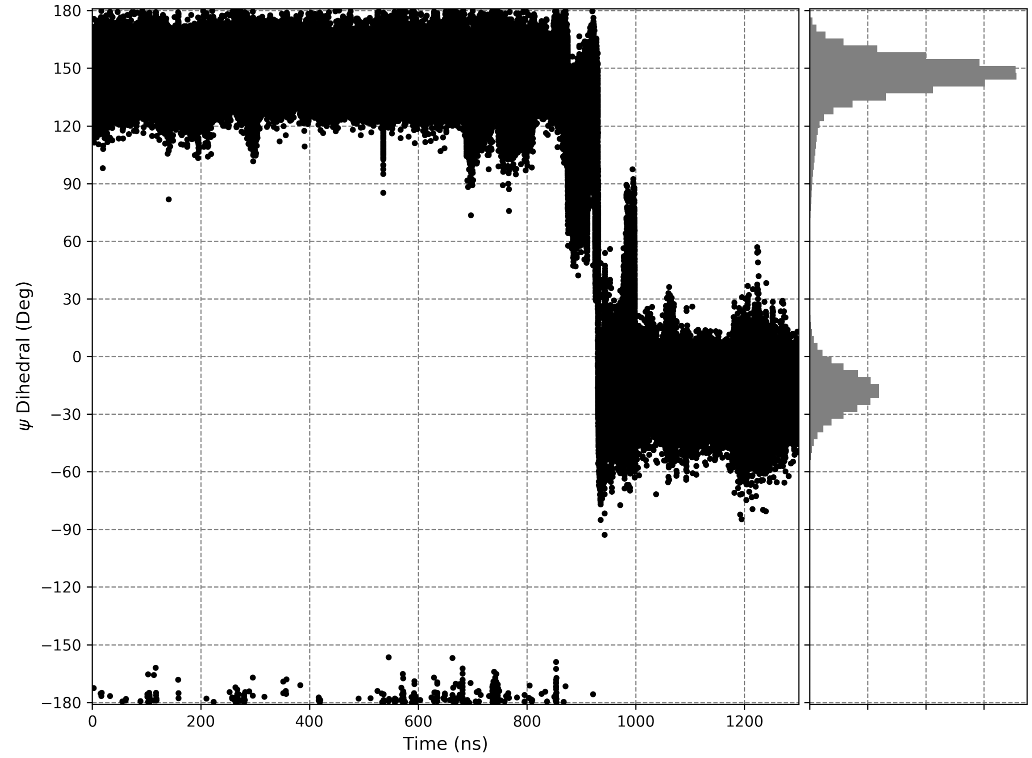


**Fig 4.** $\psi$ dihedral of Thr245 in the unbiased Apo simulation. The "open" to "closed" conformational change in L$\beta$3$\beta$4 occurs at approximately 900 ns.

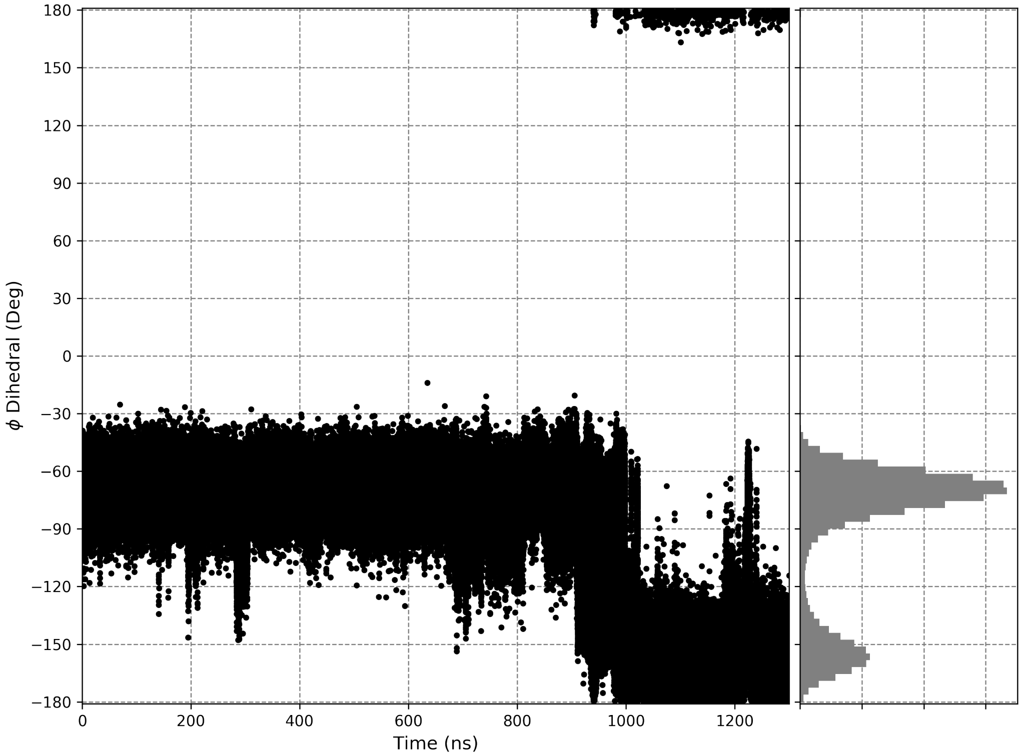


**Fig 5.** $\phi$ dihedral of Thr246 in the unbiased Apo simulation. The "open" to "closed" conformational change in L$\beta$3$\beta$4 occurs at approximately 900 ns.

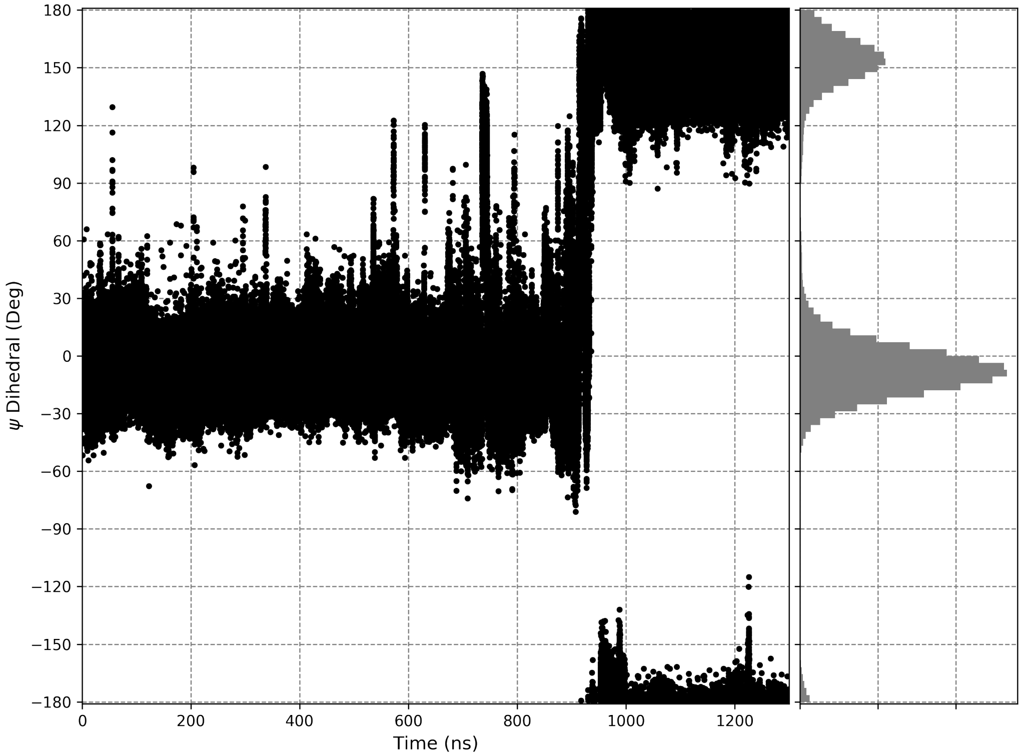


**Fig 6.** $\psi$ dihedral of Thr246 in the unbiased Apo simulation. The "open" to "closed" conformational change in L$\beta$3$\beta$4 occurs at approximately 900 ns.
